# Mapping bacterial bile amidation across the bile salt hydrolase family

**DOI:** 10.64898/2026.11.01.763480

**Authors:** Anna E. Skrip, Swati Shyam Prasad, Igor H. Wierzbicki, Vincent Charron-Lamoureux, Hunter Caleb Trithart, David Pattynama, Victoria Deleray, Carlynda Lee, Kyle Vitalli, Yuanhao Liu, Jasmine Zemlin, An Nguyen, Eden Lee, Lee R. Hagey, David J. Gonzalez, Pieter C. Dorrestein, Ipsita Mohanty, April L. Lukowski

**Affiliations:** Department of Biochemistry and Molecular Biophysics, University of California San Diego, La Jolla, CA, USA; Department of Veterinary and Biomedical Sciences, The Pennsylvania State University, University Park, PA, USA; Department of Pharmacology, University of California San Diego, La Jolla, CA, USA; Skaggs School of Pharmacy and Pharmaceutical Sciences, University of California San Diego, La Jolla, CA, USA; Center for Microbiome Innovation, University of California San Diego, La Jolla, CA, USA; Department of Medicine, University of California San Diego, La Jolla, CA, USA; Collaborative Mass Spectrometry Innovation Center, Skaggs School of Pharmacy and Pharmaceutical Sciences, University of California San Diego, La Jolla, CA, USA; Department of Nutritional Sciences, The Pennsylvania State University, University Park, PA, USA; Center for Marine Biotechnology and Biomedicine, Scripps Institution of Oceanography, University of California San Diego, La Jolla, CA, USA

## Abstract

Evaluating enzyme activity and specificity in complex microbiomes is challenging. The recent discovery that bacterial bile salt hydrolases (BSHs) also function as transaminases in the human gut microbiome highlights what remains to be disentangled, even within one of the most extensively studied microbiomes. BSHs as transaminases can conjugate bile acids with bioavailable amines to produce bacterial bile amidates (BBAs). The implications of BBAs for human health are rapidly evolving, but our understanding of which BSHs produce which specific BBAs remains limited. Here, we connect BSH sequence and function by evaluating the transaminase activities of 115 BSHs and reveal distinct amine specificities using molecular networking. Additionally, we identify enzymatically generated BBAs distributed across life-stage, tissue-, and disease-specific human samples by mining public metabolomics repositories, highlighting the previously unrecognized catalytic diversity of BSHs and their potential relevance to human health.

**Graphical Abstract:** 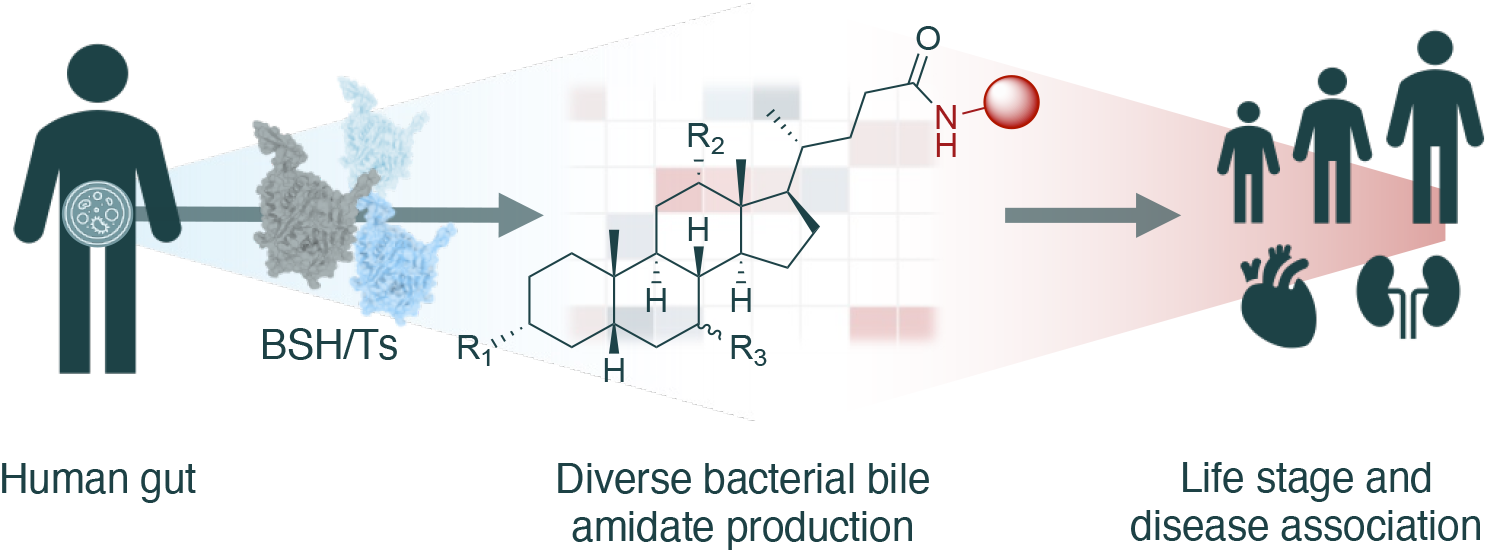

## Main

Enzyme chemistry at the human-microbe interface poses several challenges for study due to the complexity of both the human microbiome and its associated metabolome. The N-terminal nucleophile (Ntn) hydrolase superfamily, which is a diverse family that catalyzes transamination and (thio)ester and amide bond hydrolysis reactions on a variety of substrates, operates at this interface.^1,2^ The family is named for its requirement of a free N-terminal nucleophile for activity, commonly a cysteine, serine, or threonine, where the terminal methionine residue is removed autocatalytically^1,3–6^ or by a methionine aminopeptidase^7^. Members of this superfamily include bile salt hydrolases (BSHs),^1,8^ which are known for their roles as gatekeeper enzymes in the human gut microbiome that regulate the cycling of human-produced bile acids and thus represent a key biochemical link between host and microbe. BSHs were first discovered to hydrolyze the amide bond in bile acids conjugated with taurine or glycine^9^ but have recently been shown to exhibit an expanded scope of reactivity by catalyzing transamination reactions to generate bacterial bile amidates (BBAs) from amino acids and other amines^10,11^ (**Figure 1A**). Studies thus far have focused on the transamination potential of only a small subset of enzymes not representative of the BSH sequence diversity, and the catalytic capacity of the family for producing specific BBAs is consequently not well understood.

**Figure 1.**
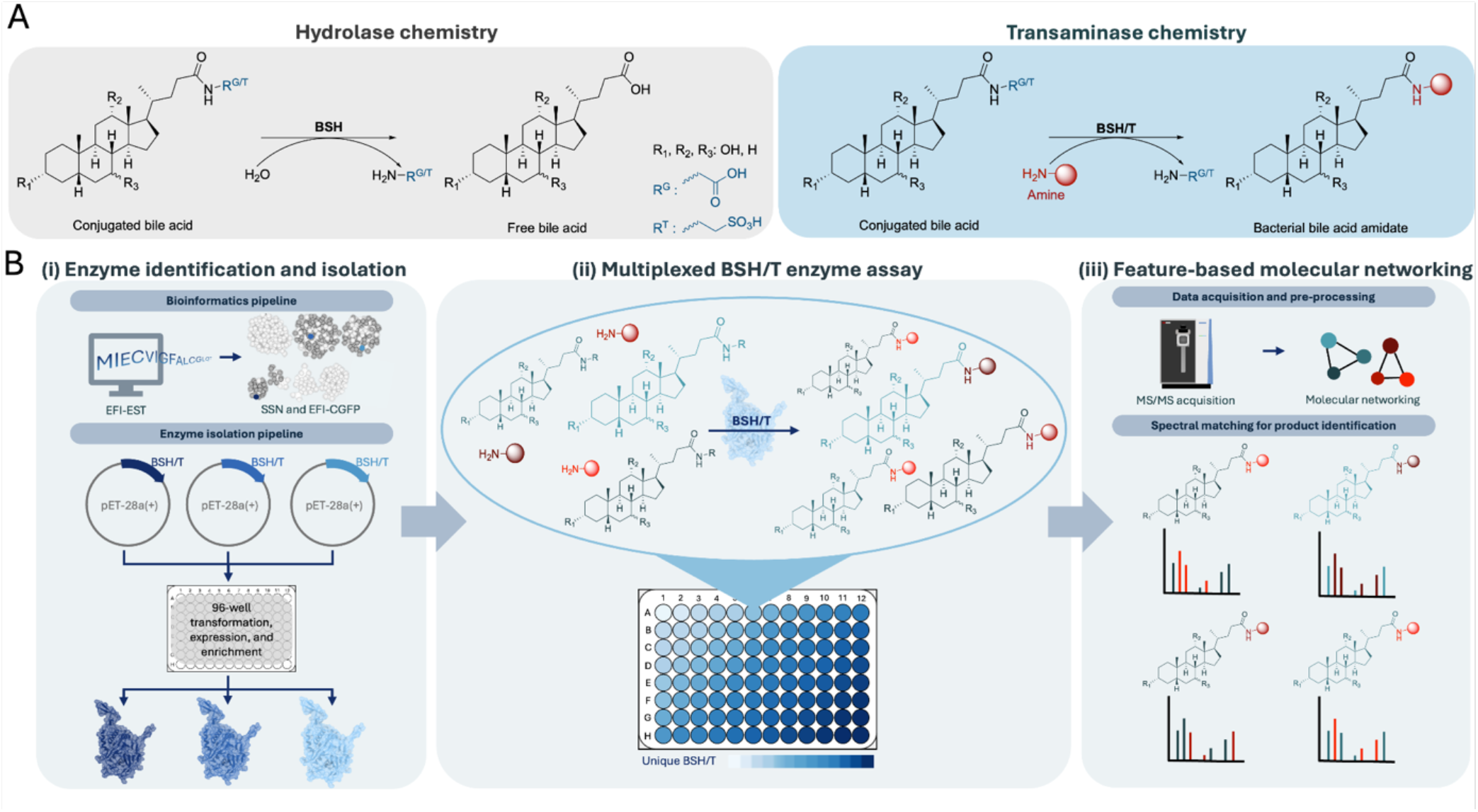
Motivation and overview of workflows. A) Comparison of BSH reactions. B) (i) BSHs were identified via the Enzyme Function Initiative Enzyme Similarity Tool (EFI-EST) and Computationally Guided Functional Profiling (CGFP), heterologously expressed in *E. coli*, and enriched in a 96-well plate format. (ii) Multiplexed assays containing substrate pools were performed in 96-well plate format with each well containing a unique BSH. (iii) Assays were analyzed by LC-MS/MS and data analysis was performed with Feature-Based Molecular Networking and spectral matching against GNPS libraries.

Thousands of structurally diverse BBAs have been identified through untargeted liquid-chromatography tandem mass spectrometry (LC-MS/MS) metabolomics approaches^12–28^. This increasing chemical repertoire presents a mismatch between the handful of known BSH transaminase products and the approximately 2,400 unique BSHs identified in microbiome sequencing studies^29–31^. Although specific BBAs continue to be associated with human health conditions, the origins of those BBAs are routinely attributed to gut microbial species, not to specific enzymes^12,13,15,18,20,24,28,32–35^. Broadly characterizing BSH transamination substrate specificities is therefore essential for establishing connections between metabolites, enzymes, microbial taxa, and human biology, but challenging because both the bile acid and amine substrates are variable. We sought to characterize the activities of over 100 enzymes representing BSH sequence diversity, focusing on enzymes that had not been previously characterized, and using a defined substrate pool representing metabolites found in the human gut (**Figure 1B**). To analyze such reactions, we envisioned using untargeted LC-MS/MS metabolomics and feature-based molecular networking to determine product formation from pooled assays. These approaches culminated in the development of a multiplexed assay that accelerated our ability to detect transamination fingerprints for 115 BSHs as well as associate the products with human metabolite datasets across disease and life stages.

We used the Enzyme Function Initiative (EFI)^36,37^ suite of tools to assess the sequence diversity of microbial BSHs, beginning by constructing a sequence similarity network (SSN) using two InterPro^38^ families encompassing microbial BSHs (IPR029132 and IPR047711), resulting in a total of 9,312 unique sequences. Since microbial BSH protein sequences exhibit a high level of similarity, we used a high alignment score cutoff for the SSN to cluster sequences that were 75% identical or higher (**Supplementary Figure S1**). To determine which BSHs in the network were attributed to the human gut microbiome, we used the EFI Computationally Guided Functional Profiling (EFI-CGFP) tool^36^, which generates protein sequence markers based on SSN clustering and maps those markers onto metagenomes across six tissue types from data from healthy individuals deposited in the Human Microbiome Project database^39,40^ (**Supplementary Figure S2**). Notably, BSHs present in human microbiomes were predominantly detected in stool compared to other tissue types.

Many of the stool-associated BSHs formed distinct clusters in the SSN (**Supplementary Figure S1**). We chose representative sequences for further analysis using CD-HIT^41^, resulting in a collection of 94 human-associated BSH sequences. Previously characterized BSHs from *Listeria*^42^ and penicillin V acylases (PVAs) from *Listeria*^43^ and *Lysinibacillus*^44,45^ with known activities were included as controls, as well as 17 non-human-associated enzymes from fish-associated microbiomes, wastewater, and benthic marine sediment to serve as functional comparisons for possible source dependence on enzyme activity. In total, 115 enzymes from 9 microbial phyla were chosen for this study (**Figure 2**).

**Figure 2.**
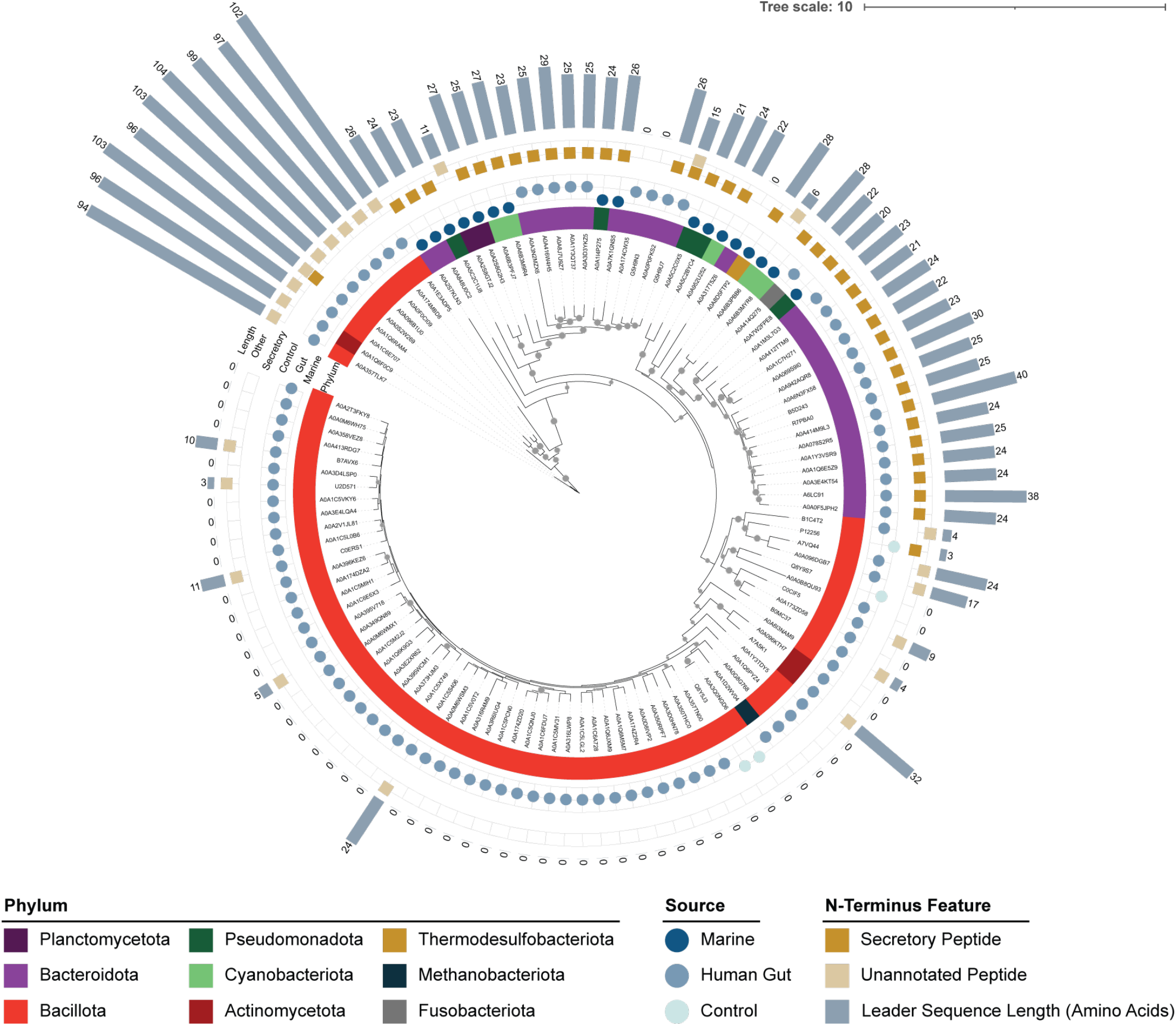
Phylogenetic distribution of selected sequences. Phylogenetic tree of selected enzymes showing phylum, source (human gut microbiome or marine microbiome-derived), presence or absence of N-terminal sequence features such as secretory signal peptides, and amino acid length of N-terminal features. The phylogenetic tree was visualized and annotated with iTOL v7^46^, with bootstraps greater than 75 (1000 iterations) indicated by grey circles at nodes.

Phylogenetic analysis revealed two distinct clades of Bacillota protein sequences, which apparently separated based on the presence or absence of N-terminal leader sequences up to 104 amino acids in length (red, **Figure 2**). Although shorter (~20-30 amino acids) leader sequences signaling transport to the periplasm have been previously reported in Ntn-hydrolases from Gram-negative microbes (i.e., Bacteroidota, purple in **Figure 2**), such long leader sequences have not been reported in the Gram-positive Bacillota BSHs^47^. By analysis with SignalP^48^ and DeepTMHMM^49^, the vast majority of the Bacteroidota sequences and Gram-negative marine-sourced sequences were predicted to contain secretory signal peptides (**Figure 2, Supplementary Figure S3**). Interestingly, alignment of the >100 amino acid long Bacillota leader sequences revealed that the first 20-30 amino acids showed similarity to Bacteroidota secretory peptides, but the remainder of the leader sequence exhibited transmembrane features (**Supplementary Figure S3**). Curious as to whether these features affected enzyme function, we chose to retain all leader sequences in this study.

To obtain enzymes for activity assays in a streamlined manner, we heterologously expressed all 115 enzymes in *Escherichia coli* in 96-well plates and used 96-well Ni-NTA and desalting plates for enrichment. The presence of each protein was confirmed by SDS-PAGE (**Supplementary Figure S4**) and mass spectrometry-based proteomics. Rather than testing combinations of substrates individually, we combined 19 bile acids, including glycocholic acids (GCA) and taurocholic acids (TCA), and 35 amines, including amino acids, polyamines, and neurotransmitters such as dopamine and tryptamine, in a substrate pool (**Supplementary Figure S5, Supplementary Figure S6**). Since each reaction had the potential to yield up to 665 unique products, we used untargeted LC-MS/MS analysis coupled with feature-based molecular networking and GNPS2^50,51^ and spectral alignment to bile acid-specific libraries to detect BBA products. Spectral library matching followed by MassQL-based bile acid filtering^12^ yielded 58 matches, which were selected for downstream analysis.

To assess transamination preferences, we compared BBA product formation across 115 enzymes by plotting mean peak areas and performing hierarchical clustering (**Figure 3**). Of these, 92 enzymes (80.0%) exhibited detectable signals for at least one BBA product, while 40 enzymes (34.8%) produced at least one BBA meeting the stringent reproducibility criteria (CV < 0.4 and log_10_ mean peak area > 1). The heatmap shows the 58 reproducibly detected BBAs, indicated by “mono-, di-, or tri-” for the level of hydroxylation on the bile steroid core, the amine detected, and the isomer number. For example, “Mono_ornithine_iso_2” indicates a monohydroxylated bile steroid conjugated to ornithine, although since ornithine has two free amines available for reaction, two isomeric products are possible and resolved by retention time. Similarly, “Di_ornithine_iso_4” is one of four possible isomers due to the inclusion of different dihydroxylated bile acid isomers in the substrate mixture. The most commonly observed BBAs were Tri-GABA (78/115; 67.8%) and Tri-2,3-diaminopropionic acid (60/115; 52.2%), followed by 3α,7α,12oxo-alanine, Tri-2-aminophenol, and Di-alanine, each produced by 50.4% (58/115) of the enzymes (**Supplementary Figure S7**).

**Figure 3.**
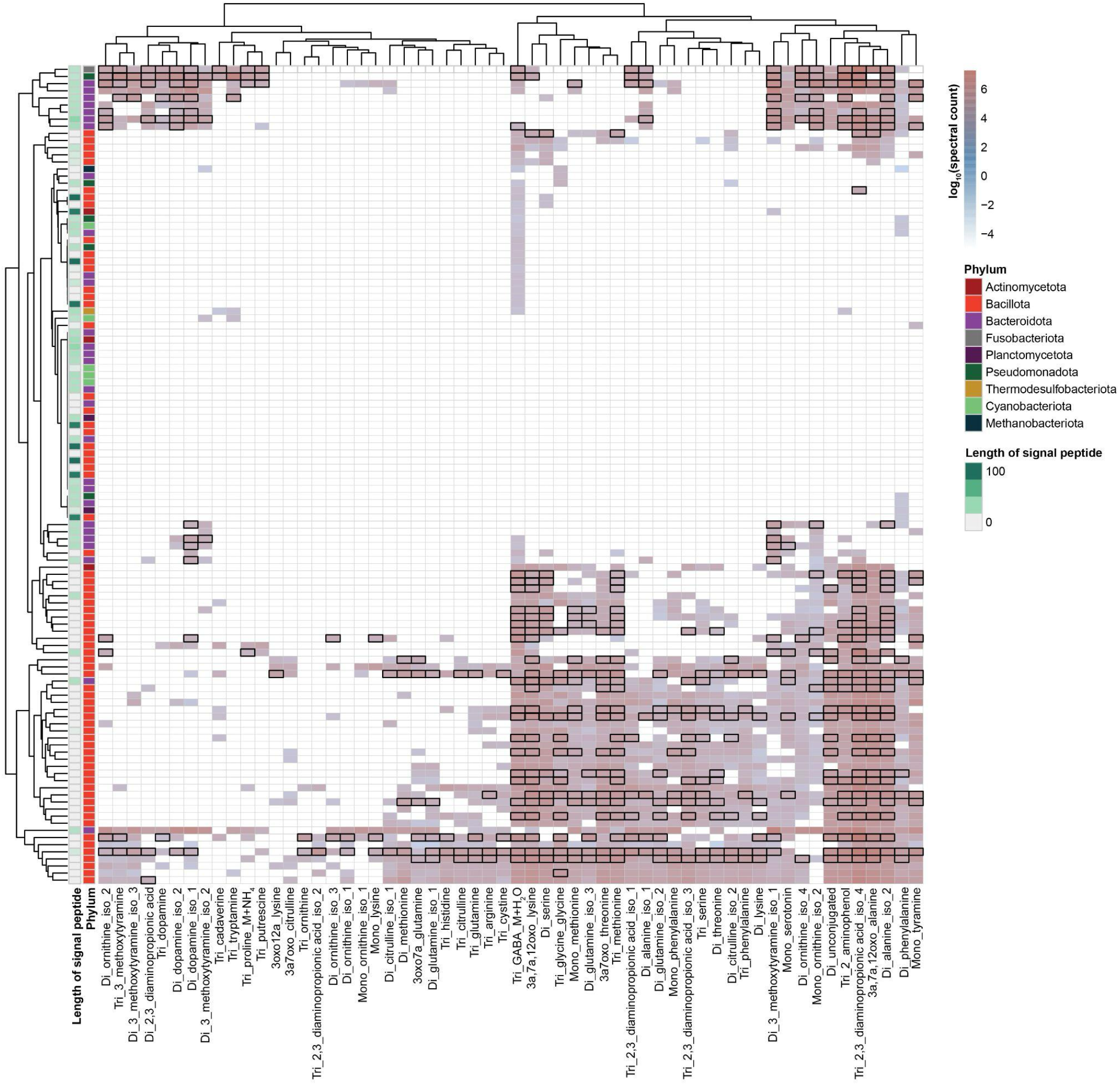
Enzyme-substrate landscape of BBAs across bacterial phyla. Heatmap of log_10_ mean peak areas for bile acid-amine conjugates across BSH enzymes in triplicate. Rows are enzymes colored by phylum and annotated with signal peptide length; columns are BBA products. Nomenclature for products are Mono-, Di-, or Tri-, representing the bile acid core followed by the amine added, and the numerical ID represents that the BBA has more than one isomer detected in the assays that cannot be structurally resolved further. Black outlines mark robust enzyme-substrate pairs (CV < 0.4, log_10_ mean > 1). White indicates no detectable signal.

Several clusters of high spectral abundance were observed throughout the heatmap, indicating groups of enzymes that efficiently catalyzed the formation of common subsets of BBAs. One subgroup of Bacillota-derived enzymes (bottom section of **Figure 3**) exhibited relatively broad substrate promiscuity, suggesting high activity in producing a diverse set of BBAs. Notably, these were the Bacillota sequences lacking signal peptide sequences, and the Bacillota enzymes harboring long signal peptides were inactive in this assay. Comparatively, a group of Bacteroidota BSHs (top section of **Figure 3**) produced high spectral counts of BBAs containing amines that were less commonly detected in the broader collection of enzymes, namely methoxytyramine, tyramine, serotonin, alanine, and 2-aminophenol. Additionally, several BBAs formed isolated clusters in the heatmap resulting from processing by a small subset of enzymes, indicating that certain specialized BSHs have rare specificities toward structurally uncommon conjugates. The clustering pattern also indicates that the conjugated amine and the bile steroid core jointly guide substrate preference, rather than either substrate driving transamination specificity alone. Overall, these data demonstrate that BSH-mediated transamination spans a continuum from highly promiscuous to highly specialized enzymes, indicating that BSHs in the gut microbial consortium partition the BBA chemical space into distinct substrate classes.

We next queried the MS/MS spectra of the enzymatically produced BBAs against more than 2 billion MS/MS spectra deposited in the public repositories GNPS/MassIVE^51^, MetaboLights^52^, and Metabolomics Workbench using fastMASST^53–55^ to determine whether these products are observed beyond our experimental system (**Figure 4A**). To place these spectral matches into biological context, we integrated repository metadata using PanReDU^55^, a controlled-vocabulary framework that harmonizes sample annotations across public metabolomics datasets, focusing on disease associations and life-stage metadata. We detected MS/MS spectra for 22 of the 58 BBAs identified in our enzyme assays in at least one disease-associated metabolomics study (**Figure 4B**), demonstrating that many of these conjugates or close structural isomers occur in diverse biological settings.

**Figure 4.**
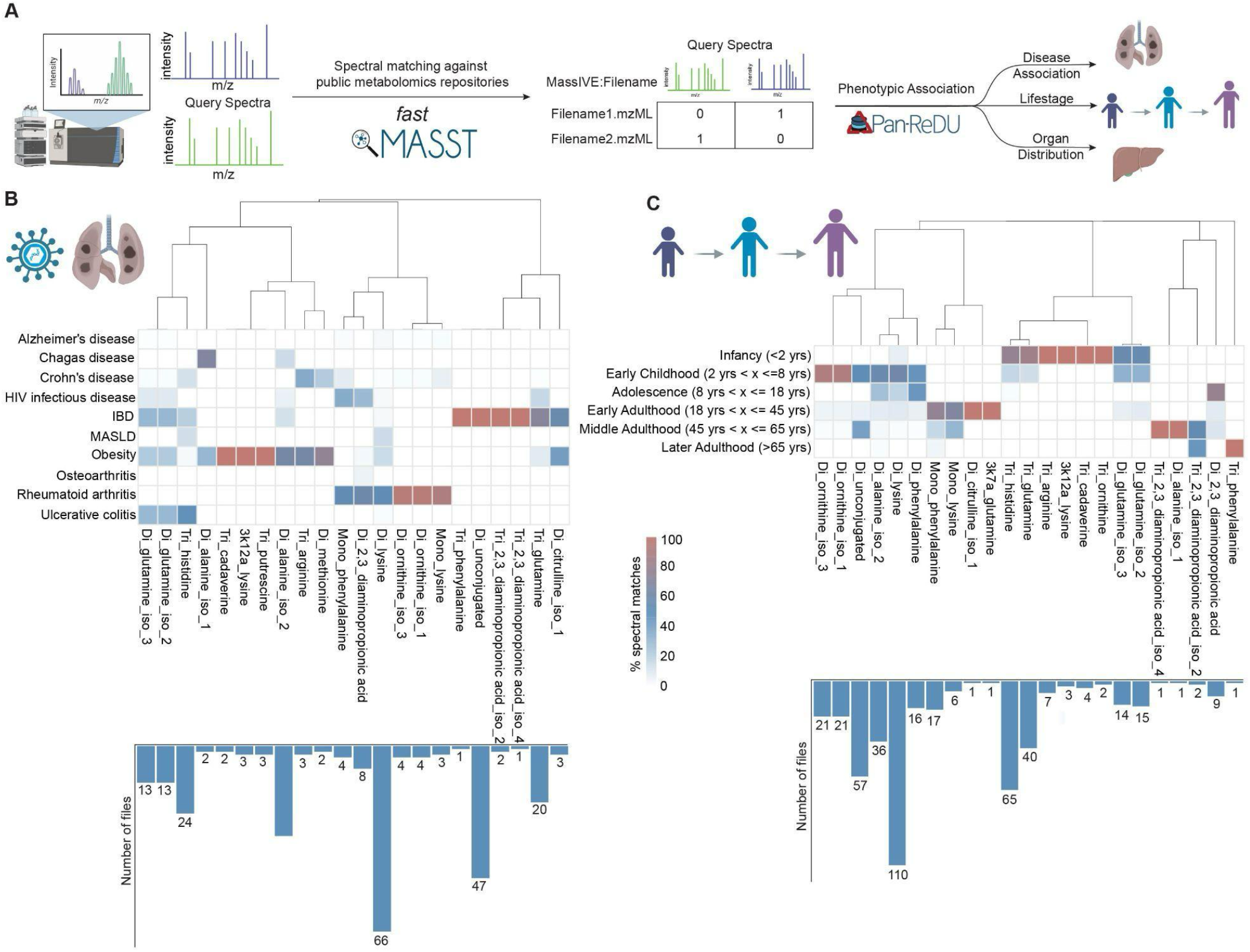
Repository-scale phenotypic associations. A) Schematic overview of the workflow used to contextualize bile acid conjugates across public metabolomics repositories. B) Heatmap showing the percentage of spectral matches for bile acid conjugates across human disease-associated datasets. C) Heatmap showing the prevalence of bile acid conjugates across different human life stages, including infancy, childhood, adolescence, adulthood, and later adulthood. In both B and C, bar plots below indicate the total number of files containing spectral matches for each metabolite.

Among all conjugates, dihydroxy-lysine bile acid was the most frequently detected, yielding 66 spectral matches across public datasets, and occurred most frequently in rheumatoid arthritis datasets, followed by obesity and metabolic dysfunction-associated steatotic liver disease (MASLD) datasets, suggesting that this conjugate is detected across multiple inflammatory and metabolic disease contexts. Additionally, we detected putative trihydroxylated bile acid-2,3-diaminopropionic acid (DAP) conjugates that, to our knowledge, have not previously been reported. This conjugate matched to IBD datasets at low frequency. The dihydroxylated-DAP, in which deoxycholyl-DAP was previously reported^56^, was not detected in IBD, but was detected in rheumatoid arthritis and HIV datasets, suggesting that the bile steroid core may contribute to disease-associated distribution patterns. Across all health categories examined, IBD, rheumatoid arthritis, and obesity contained the greatest BBA diversity, indicating that bile acid amidation may be particularly prominent in these disease-associated metabolomes.

Life-stage analysis also revealed distinct patterns of BBA occurrence (**Figure 4C**). Conjugates containing polyamines and related amino acid metabolites, including arginine, ornithine, and cadaverine, were detected almost exclusively in samples from infants (<2 years), suggesting that these BBAs may reflect age-dependent microbial metabolic activities during early life. In contrast, trihydroxy-phenylalanine bile acid was detected only in later adulthood (>65 years). Collectively, these analyses demonstrate that both established and previously undescribed BBAs are detectable across independent human metabolomics studies and exhibit distinct disease- and age-associated distributions, providing evidence that these microbial bile acid conjugates are biologically relevant features of the human metabolome.

In summary, we developed an efficient multiplexed assay to characterize the transamination potential of 115 BSHs using a biologically relevant substrate pool. By incorporating feature-based molecular networking, we facilitated product identification and searchability in public metabolomics repositories, resulting in the association of novel biocatalytic reaction products from microbial enzymes to human-sourced metabolite extracts. Through these analyses, we observed that members of the BSH family in the human gut range from promiscuous to selective for transamination. Although this is the largest BSH-substrate screen performed to date, it remains limited because not all described biological amines or bile amidates could be tested; however, we anticipate that our approach is both scalable and adaptable. As the biological roles of specific BBAs continue to be elucidated, resolving which enzymes make which products will ultimately be essential to understanding how BBAs shape, and are shaped by, host-microbiome interactions, as well as their broader impacts on human health.

## Associated Content

All proteomics data (.raw file and metadata) are publicly available online at MassIVE Repository (https://massive.ucsd.edu) under identifier: MSV000103172. Untargeted LC-MS/MS data is deposited to GNPS/MassIVE under the accession MSV000097665. Code for Figure 3 and Supplementary Figure S7 can be accessed here: https://github.com/mohantyipsita/2026_BSH_substrate_screening. Code for Figure 4 can be accessed here: https://github.com/ssp5682-create/BSH_study_code. Additional supplementary data files associated with this work, including protein sequences, proteomics data summaries, mass spectral data tables, and spectral matching results, are available at Zenodo with DOI: 10.5281/zenodo.23814337 Methods and Supplementary Figures are provided in: Supporting Information PDF

## Author Information

## Acknowledgements

This work was supported by the Molecular Biophysics Training Grant, NIH Grant T32 GM139795 (to A. E. S.), Health Sciences Collaborative Center for Proteomics (D.J.G founding director), startup funds from Penn State to I.M., R01 DK136117 (P.C.D) and startup funds from UC San Diego to A.L.L.

