## Supporting Information for "Mapping bacterial bile amidation across the bile salt hydrolase family"

### Table of Contents

**I. Bioinformatics…………………………………………………………………………. 3**
Enzyme Candidate Selection…………………………………………………………….. 3
Phylogenetic Tree Construction………………………………………………………… 3
Figure S1. Sequence Similarity Network……………………………………… 4
Figure S2. Metagenome Marker Abundance………………………………….. 5
Figure S3. Sequence Alignment of Leader Peptide Sequences…………….. 6
**II. Enzyme Expression and Purification……………………………………………... 7**
General Considerations…………………………………………………………………... 7
Selection of candidate bile acid amidates (BBAs) …………………………………... 15
Repository-scale spectral searching using fastMASST……………………………... 16
Mapping BBA spectral matches to public metabolomics metadata…………………. 16
Figure S7. Frequency of the most commonly detected BBAs across the enzyme screen……………………………………………………………….. 17
**V. Proteomics…………………………………………………………………………… 18**
Sample Preparation……………………………………………………………………….. 18
Data Acquisition…………………………………………………………………………… 19
Data Analysis……………………………………………………………………………… 19
Data and code availability……………………………………………………………….. 20

### I. Bioinformatics

**Enzyme candidate selection.** The InterPro families IPR029132 (choloylglycine hydrolase, NAAA C-terminal) and IPR047711 (conjugated bile acid hydrolase) were chosen for analysis using the Enzyme Function Initiative Enzyme Similarity Tool (EFI-EST), resulting in a total of 11,203 sequences after duplicates were removed. Fragmented sequences were removed, resulting in a total of 9,312 sequences. The majority of the sequences were of bacterial origin ranging from 300-390 amino acids in length. Several alignment scores (ranging 115-170) were used to generate the SSN to determine optimal clustering of sequences, ultimately using an alignment score of 170 (approx. 75% sequence identity in clusters). This network was further analyzed by the EFI Chemically-Guided Functional Profiling (EFI-CGFP) tool using a default CD-HIT sequence identity of 85% and analyzed against all metagenome marker types (anterior nares, buccal mucosa, supragingival plaque, tongue dorsum, stool, and posterior fornix) in a collection of 380 metagenomes derived from healthy adult males and females deposited in the Human Microbiome Project database. The vast majority of BSH sequence markers (1501) were identified in stool. SSN clusters containing two or more human-associated markers were analyzed by CD-HIT at a 70% cutoff to statistically determine the representative sequence from each cluster. In cases where only part of a cluster was identified in human metagenomes, only the human metagenome-associated sequences were used in the CD-HIT analysis. From this analysis, 94 sequences were selected for study.

Control BSH sequences were chosen based on literature precedence for hydrolase or acyltransferase activity. Non-human-associated sequences were selected from SSN clusters lacking human-associated markers to emphasize the functional diversity of the enzyme family across distinct environments. Marine bacteria were selected based on host associations (fish, marine algae, benthic environments) and their presence in high-quality genomic data, yielding 17 additional enzyme sequences.

**Phylogenetic tree construction.** Selected sequences described above were aligned using MUSCLE[64] (v3.8) and an unrooted maximum likelihood phylogenetic tree was created using IQ-TREE[65] (v2.1.4-beta COVID-edition) in automated model selection mode with the best substitution model found to be LG+I+G4. The consensus tree was constructed from 1000 bootstrap trees and visualized and annotated using iTOL v7[50]. The annotated tree can be viewed interactively at the following link: https://itol.embl.de/shared/Lukowski_Lab

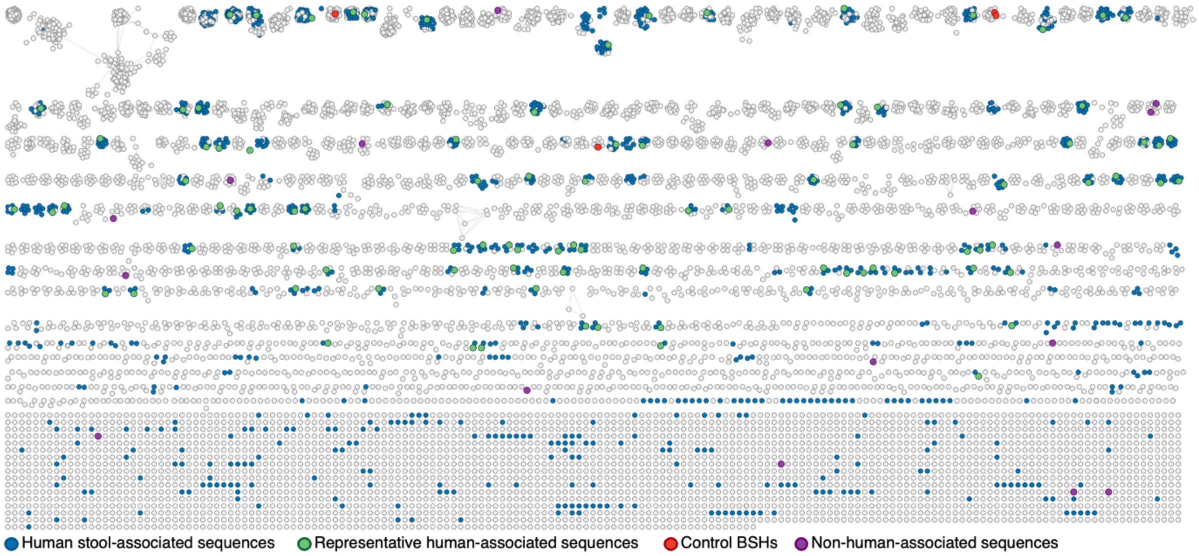

**Supplementary Figure S1. Sequence Similarity Network (SSN).** Microbial sequences from IPR029132 and IPR047711 are represented in an SSN produced at an alignment score of 170. Each node corresponds to one individual protein sequence. Nodes in blue indicate protein sequences detected in human fecal sample microbiome data, nodes highlighted in green represent the human-associated sequences chosen for analysis in this study, with one representative per human stool-associated cluster shown; nodes in red represent control BSHs, and nodes in purple represent non-human-associated sequences.

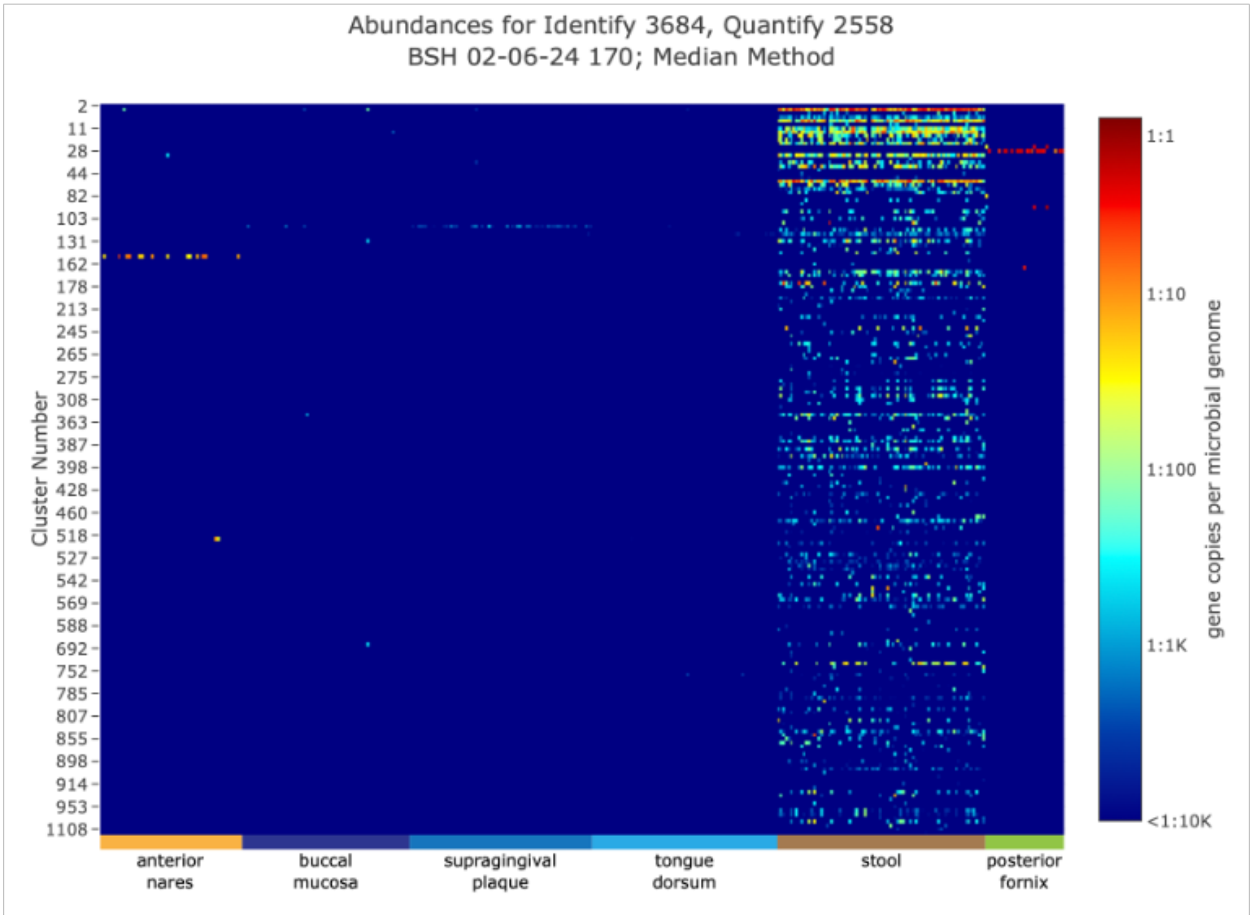

**Supplementary Figure S2. Metagenome marker abundance.** Gene copies identified per microbial genome for markers generated by the EFI-CGFP tool based on clusters defined by the EFI-EST ColorSSN tool (y-axis). Of the six human tissue types tested, BSH sequences were most abundant in stool.

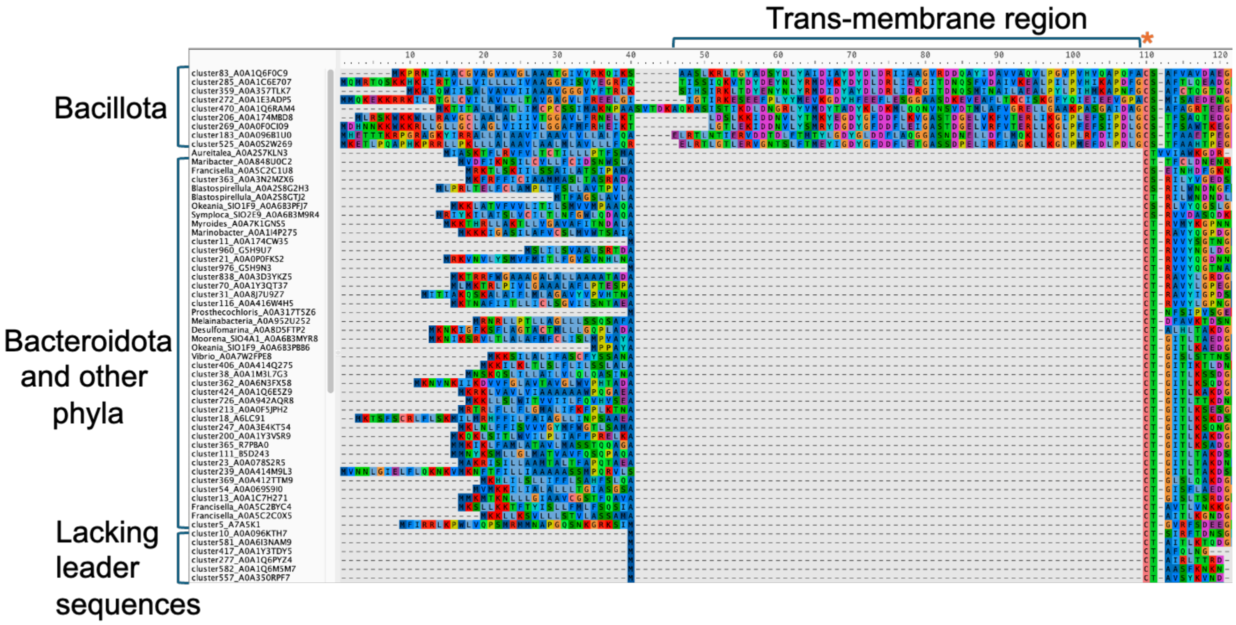

**Supplementary Figure S3. Sequence alignment of leader peptide sequences.** Screenshot of a portion of the sequence alignment of BSHs in this study generated with MUSCLE v3.8 and visualized with Aliview v1.28. The top nine sequences are the Bacillota with long (>100 amino acids) leader peptides and the lower sequences are the Bacteroidota sequences and examples from other phyla with 20-30 amino acid length peptides, demonstrating the similarity between these peptides and the first 20-30 amino acids in the Bacillota sequences. The cysteine residue highlighted in orange toward the right is the conserved catalytic cysteine.

### II. Enzyme Expression and Purification

**General considerations.** All enzyme sequences were codon-optimized for expression in *E. coli,* synthesized by Twist Bioscience, and inserted into pET28a(+) vectors between *Nco*I and *Xho*I to afford C-terminal 6x-polyhistidine-tagged proteins. All bile acid and amine substrates were sourced commercially.

**BSH expression in 96-well plate format.** The BSH sequences selected from the sequence similarity network were codon-optimized for *Escherichia coli* and inserted into pET28a(+) vectors between *Nco*I and *Xho*I restriction sites with a kanamycin resistance gene and a C-terminal 6x-Histidine tag (synthesized by Twist Biosciences). Constructs were ordered in a 96-well plate format and transformed into chemically competent *Escherichia coli* BL21(DE3) cells using heat-shock transformation. Transformations were recovered in 250 µL of LB media for 60 minutes, and 50 µL were transferred to 500 µL of Luria-Bertani (LB) media with 50 µg/mL kanamycin. This starter culture was incubated overnight at 37 °C, 850 rpm. An expression culture was prepared by inoculating 500 µL LB media containing 50 ug/mL kanamycin with 20 µL of starter culture. The expression culture was incubated for 2 h at 37 °C, 850 rpm until the optical density at 600 nm (OD600) was between 0.6-0.8, then expression was induced with Isopropyl β-D-1-thiogalactopyranoside (IPTG) to a final concentration of 1 mM. Induced cultures were incubated overnight at 18 °C, 850 rpm. Cultures were pelleted by centrifugation for 30 min at 4 °C, 4000 x *g*, and the supernatant was discarded. Cell pellets were stored at –80 °C until purification.

**BSH enrichment in 96-well plate format.** Cell pellets were thawed on ice and resuspended in 400 µL lysis buffer (50 mM Tris-HCl, pH 7.5 500 mM NaCl, 10% glycerol, 10 mM imidazole supplemented with 1 mg/mL lysozyme, 0.5 mg/mL polymyxin B, 10 µg/mL DNase I, 1 mM PMSF) and incubated for 2 hours at 30 °C, 950 rpm. Lysates were pelleted at 7000 xg, 30 mins, 4 °C, and the crude supernatant was transferred to a His MultiTrap plate (Cytiva) that was pre-equilibrated with lysis buffer. Bound proteins were washed twice with 100 µL of lysis buffer. His-tagged proteins were eluted with 200 µL of elution buffer (50 mM Tris-HCl, pH 7.5 500 mM NaCl, 10% glycerol, 500 mM imidazole) and transferred to a 7K molecular weight cutoff (MWCO) Zeba Spin Desalting Plate (ThermoFisher) that was pre-equilibrated with desalting buffer (50 mM Tris-HCl, pH 7.5 300 mM NaCl, 10% glycerol). Proteins were eluted in 200 µL of desalting buffer by centrifugation at 100 x *g*, 4 °C, 2 mins, and stored at –80 C.

| Lane | Accession ID | MW (kDa) | MW – SigPep (kDa) | Sigpep detected | Activity |
|---|---|---|---|---|---|
| 1 | Ladder | - | - | - | - |
| 2 | A0A0P0FKS2 | 41.9 | 38.9 | No | No |
| 3 | A0A069S9I0 | 42.0 | 39.8 | No | No |
| 4 | B5D243 | 42.3 | 39.6 | No | No |
| 5 | A0A174Z2R4 | 39.7 | - | - | Yes |
| 6 | A0A1C5L0B6 | 39.0 | - | - | Yes |
| 7 | A0A2V1JL81 | 39.0 | - | - | No |
| 8 | A0A1C5MV31 | 39.6 | - | - | Yes |
| 9 | A0A1Q6E5Z9 | 41.5 | 39.1 | No | No |
| 10 | A0A1C5X749 | 39.3 | - | - | No |
| 11 | A0A1Q6M5M7 | 39.1 | - | - | No |

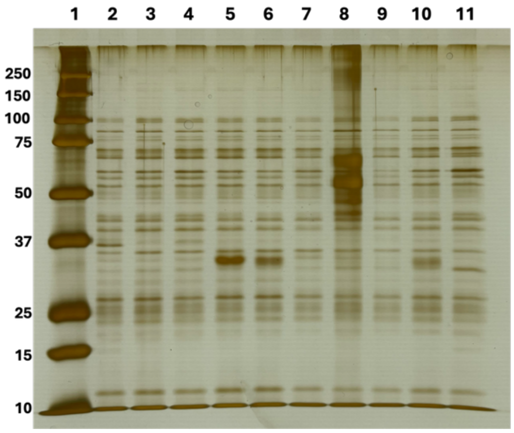

| lane | sample | MW (kDa) | MW – SigPep (kDa) | SigPep detected | Activity |
|---|---|---|---|---|---|
| 1 | A7A5K1 | 40.8 | 37.0 | Yes (6->37) | No |
| 2 | A0A078S2R5 | 41.9 | 39.4 | No | Robust |
| 3 | A0A1C6E6X3 | 38.7 | - | - | Robust |
| 4 | Ladder | - | - | - | - |
| 5 | A0A416W4H5 | 40.8 | 38.2 | No | Yes |
| 6 | A0A1C5QNJ0 | 38.9 | - | - | Robust |
| 7 | A0A395WCM1 | 38.6 | - | - | Yes |
| 8 | A0A1Q6K9G3 | 39.0 | - | - | Robust |
| 9 | A0A357TLK7 | 42.9 | 32.3 | Yes (38->111) | Yes |
| 10 | A0A0D8IVP2 | 38.4 | - | - | Robust |
| 11 | A0A316R4M9 | 38.1 | - | - | Yes |
| 12 | B7AVX6 | 39.9 | 38.8 | Yes (1->16) | Yes |
| 13 | A0A3D0HN78 | 39.0 | - | - | No |

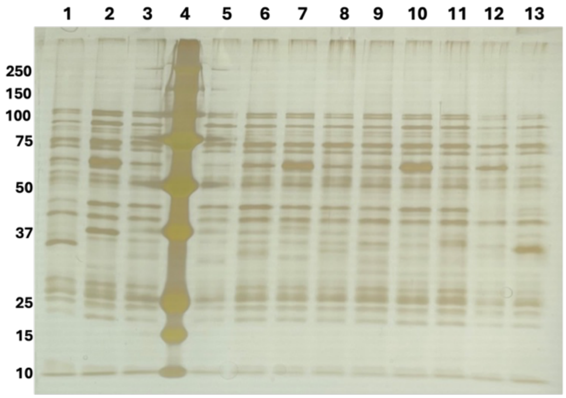

| lane | sample | MW (kDa) | MW – SigPep (kDa) | SigPep detected | Activity |
|---|---|---|---|---|---|
| 1 | U2D571 | 39.4 | 39.0 | Yes (1->10) | Yes |
| 2 | Ladder | - | - | - | - |
| 3 | A0A8J7U9Z7 | 41.4 | 38.4 | No | No |
| 4 | A0A174DZA2 | 38.4 | - | - | Robust |
| 5 | A0A1C5LGL2 | 39.0 | - | - | Robust |
| 6 | A0A174ZD20 | 39.1 | - | - | Robust |
| 7 | A0A414M9L3 | 44.2 | 39.8 | Yes (38->49) | Robust |
| 8 | A0A413RDG7 | 38.9 | - | - | Robust |
| 9 | A0A6N3FX58 | 42.5 | 39.3 | No | Robust |
| 10 | A0A1C5M2J2 | 39.3 | - | - | Yes |
| 11 | A0A395V718 | 38.8 | - | - | Robust |
| 12 | A0A173ZD58 | 39.1 | - | - | Yes |

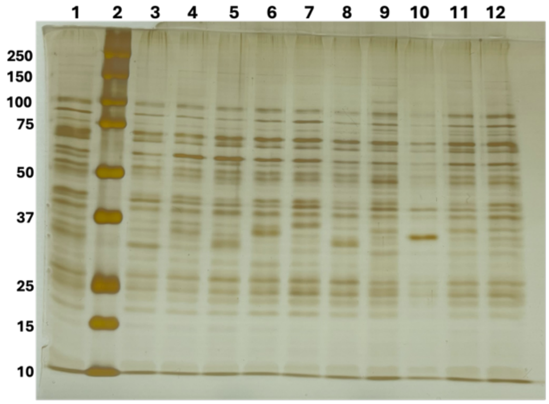

| lane | sample | MW (kDa) | MW – SigPep (kDa) | SigPep detected | Activity |
|---|---|---|---|---|---|
| 1 | A0A1C5S406 | 41.6 | 38.6 | Yes (1->31) | Yes |
| 2 | A0A0B8QU93 | 38.7 | - | - | Yes |
| 3 | Ladder | - | - | - | - |
| 4 | A0A1Y3QT37 | 41.1 | 38.5 | Yes (5->28) | Robust |
| 5 | C0ERS1 | 40.1 | 39.0 | No | Robust |
| 6 | A0A096B1U0 | 41.7 | 30.1 | Yes (44->50) | Yes |
| 7 | A0A096DGB7 | 42.8 | 41.0 | No | Yes |
| 8 | A0A0F0CI09 | 42.6 | 31.4 | No | Yes |
| 9 | A0A3N2MZX6 | 40.5 | 37.9 | No | Robust |
| 10 | A7VQ44 | 44.7 | 41.8 | Yes (3->36) | Robust |
| 11 | A0A350RPF7 | 39.1 | - | - | Robust |

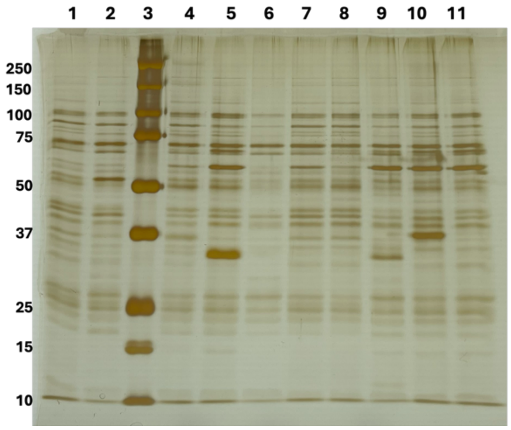

| lane | sample | MW (kDa) | MW – SigPep (kDa) | SigPep detected | Activity |
|---|---|---|---|---|---|
| 1 | A0A096KTH7 | 37.9 | - | - | Yes |
| 2 | A0A1M3L7G3 | 42.6 | 40.2 | No | Robust |
| 3 | A0A1Q6F0C9 | 41.1 | 31.0 | Yes (5->28) | Yes |
| 4 | Ladder | - | - | - | - |
| 5 | A0A1C5VKY6 | 38.7 | - | - | Robust |
| 6 | A0A1D2WV04 | 38.0 | - | - | Yes |
| 7 | A0A3E4KT54 | 42.5 | 39.8 | No | Robust |
| 8 | A0A1E3ADP5 | 41.9 | 30.4 | Yes (36->85) | No |
| 9 | R7PBA0 | 42.0 | 39.3 | No | Robust |
| 10 | A0A1C5MIH1 | 38.3 | - | - | Robust |
| 11 | A0A1Q6RAM4 | 41.6 | 31.4 | Yes (24->103) | No |
| 12 | A0A373HJM3 | 39.2 | - | - | Yes |
| 13 | A0A942AQR8 | 42.7 | 40.1 | No | Robust |

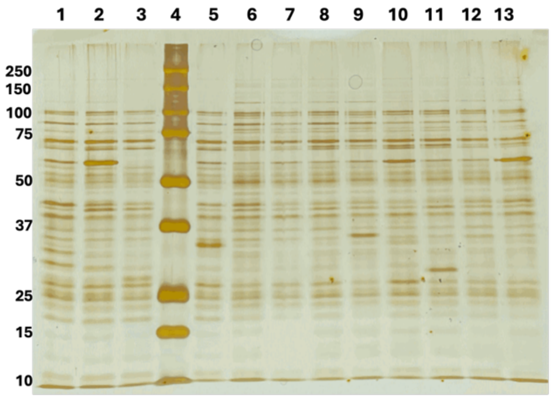

| lane | sample | MW (kDa) | MW – SigPep (kDa) | SigPep detected | Activity |
|---|---|---|---|---|---|
| 1 | Ladder | - | - | - | - |
| 2 | A0A174CW35 | 39.0 | - | - | Yes |
| 3 | A0A0M6WSM3 | 39.5 | - | - | Yes |
| 4 | A0A1C5V0T2 | 38.9 | - | - | Robust |
| 5 | A0A358VEZ8 | 38.9 | - | - | Robust |
| 6 | A0A1Y3VSR9 | 42.8 | 39.9 | Yes (25->34) | Robust |
| 7 | A0A3R6IUG4 | 38.6 | - | - | Robust |
| 8 | A0A1Q6PYZ4 | 38.3 | - | - | Robust |
| 9 | A0A412TTM9 | 41.8 | 39.3 | No | Robust |
| 10 | A0A414Q275 | 43.3 | 40.9 | No | Robust |
| 11 | B0MC37 | 39.0 | 38.5 | No | Robust |
| 12 | A0A1Q6JXM9 | 39.4 | - | - | Yes |
| 13 | A0A3E2XR62 | 38.7 | - | - | Yes |

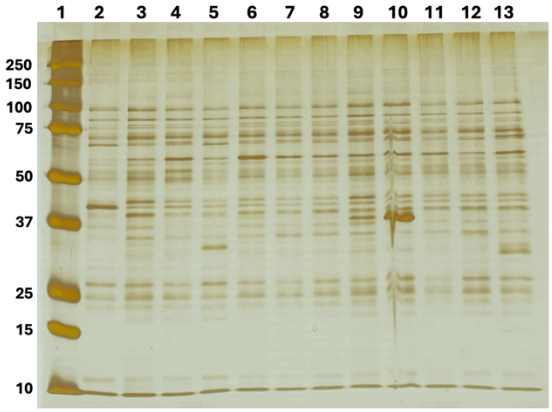

| lane | sample | MW (kDa) | MW – SigPep (kDa) | SigPep detected | Activity |
|---|---|---|---|---|---|
| 1 | A0A1C7H271 | 41.8 | 39.3 | No | Yes |
| 2 | Ladder | - | - | - | - |
| 3 | A0A1C5PCN0 | 38.4 | - | - | Robust |
| 4 | A0A0M6WMX1 | 38.6 | - | - | Robust |
| 5 | A0A0G8G768 | 39.0 | - | - | Yes |
| 6 | A0A174MBD8 | 43.1 | 32.3 | Yes (45->82) | No |
| 7 | A0A0M6WH75 | 38.9 | - | - | Yes |
| 8 | A0A1C6E707 | 44.6 | 32.7 | Yes (48->98) | No |
| 9 | A0A357TN00 | 38.8 | - | - | Robust |
| 10 | A0A1Y3TDY5 | 39.0 | - | - | Robust |
| 11 | C0CIF5 | 40.1 | 38.8 | Yes (9->18) | Yes |
| 12 | A0A350THC0 | 37.7 | - | - | Yes |
| 13 | A0A1C6FDU7 | 38.2 | - | - | No |

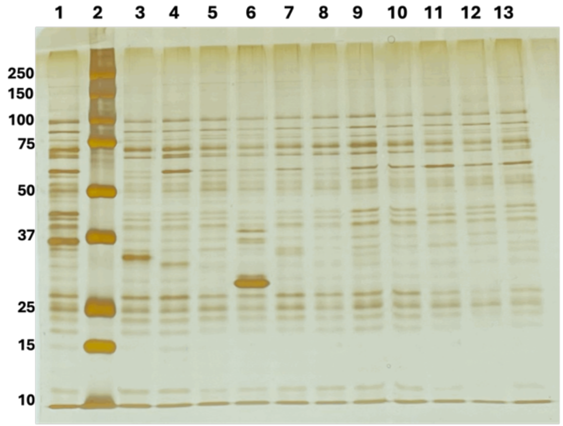

| lane | sample | MW (kDa) | MW – SigPep (kDa) | SigPep detected | Activity |
|---|---|---|---|---|---|
| 1 | A6LC91 | 44.1 | 39.8 | No | No |
| 2 | A0A2T3FKY8 | 38.8 | - | - | Yes |
| 3 | Ladder | - | - | - | - |
| 4 | A0A349QN89 | 39.4 | 38.7 | No | Yes |
| 5 | A0A1C6A728 | 40.0 | - | - | Yes |
| 6 | A0A0F5JPH2 | 42.2 | 39.3 | Yes (22->31) | Robust |
| 7 | A0A396KEZ6 | 38.7 | - | - | Robust |
| 8 | A0A3E4LQA4 | 39.5 | - | - | Robust |
| 9 | A0A316LWP8 | 39.0 | - | - | Robust |
| 10 | A0A3D4LSP0 | 38.6 | - | - | Yes |
| 11 | A0A0S2W269 | 41.7 | 30.3 | Yes (3->50) | Yes |
| 12 | A0A6I3NAM9 | 39.3 | - | - | Yes |
| 13 | A0A3D3YKZ5 | 40.8 | 38.3 | No | No |

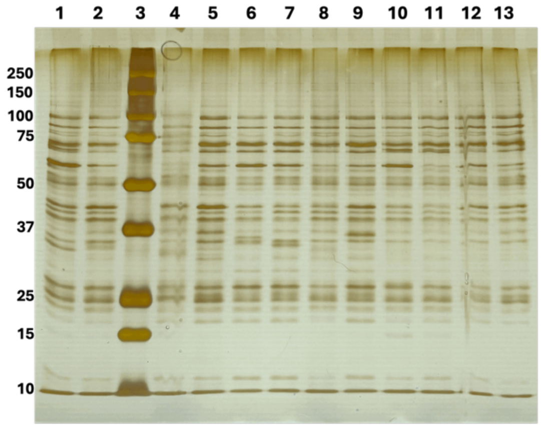

| lane | sample | MW (kDa) | MW – SigPep (kDa) | SigPep detected | Activity |
|---|---|---|---|---|---|
| 1 | P12256 | 39.8 | 39.4 | yes | Yes |
| 2 | A0A317T5Z6 | 41.1 | - | - | No |
| 3 | A0A6B3MYR8 | 44.3 | 41.2 | No | No |
| 4 | Ladder | - | - | - | - |
| 5 | A0A5C2BYC4 | 45.7 | 42.9 | No | Yes |
| 6 | A0A6B3PBB6 | 42.5 | 41.8 | Yes (2->13) | No |
| 7 | G5H9U7 | 39.7 | 38.1 | No | Yes |
| 8 | A0A848U0C2 | 43.6 | 40.8 | No | Yes |
| 9 | A0A5C2C0X5 | 44.6 | 42.4 | No | Yes |
| 10 | A0A6B3PFJ7 | 48.7 | 46.0 | No | Yes |
| 11 | A0A3Q0NGD6 | 39.1 | - | - | Robust |
| 12 | A0A2S8GTJ2 | 38.8 | 37.7 | No | Yes |
| 13 | B1C4T2 | 40.5 | 40.0 | Yes (3->21) | Yes |

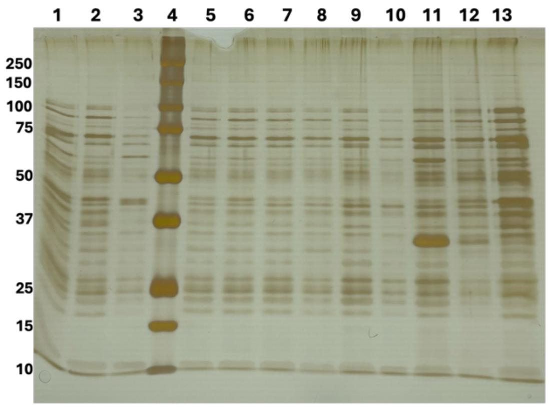

| lane | sample | MW (kDa) | MW – SigPep (kDa) | SigPep detected | Activity |
|---|---|---|---|---|---|
| 1 | G5H9N3 | 38.5 | - | - | Yes |
| 2 | Ladder | - | - | - | - |
| 3 | A0A7K1GNS5 | 41.1 | 38.3 | Yes (11->29) | No |
| 4 | A0A5C2C1U8 | 41.0 | 38.6 | No | Yes |
| 5 | A0A952U252 | 41.0 | 38.6 | No | No |
| 6 | Q8Y9S7 | 39.0 | - | - | Yes |
| 7 | A0A2S7KLN3 | 34.6 | 31.7 | No | Yes |
| 8 | A0A7W2FPE8 | 42.4 | 40.3 | No | Robust |
| 9 | A0A6B3M9R4 | 41.6 | 38.5 | No | No |
| 10 | Q8Y5J3 | 39.1 | - | - | Robust |
| 11 | A0A2S8G2H3 | 41.0 | 38.0 | No | No |
| 12 | A0A8D5FTP2 | 42.3 | 39.4 | No | Yes |
| 13 | A0A1I4P275 | 40.6 | 38.0 | No | Yes |

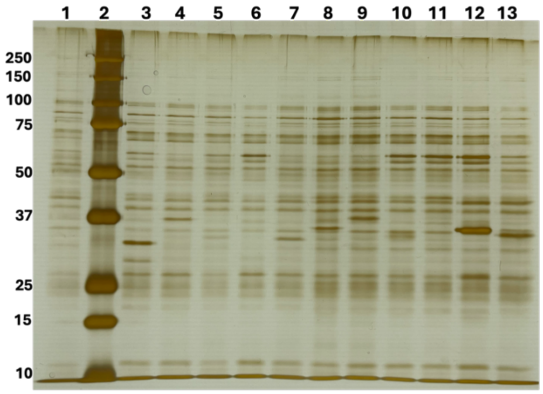

**Supplementary Figure S4. SDS-PAGE images of Ni-NTA-enriched BSHs.** After desalting, BSHs were loaded onto a 10% SDS-polyacrylamide gel. The gels were stained using a silver stain kit for band visualization. Each gel has a corresponding table of accession ID, expected molecular weights, and activity presence associated with each lane. The molecular weight ladder in kDa is on the left of each gel.

### III. Enzymatic Reactions

**Enzyme assays.** Amines (**Figure S5**) were solubilized in DMSO or Ultrapure Water to a final concentration dependent on the individual solubility of each amine (**Figure S5**). An amine master mix was prepared by mixing all amines to a final concentration of 4.2 mM for each amine in Tris-HCl, pH 4.6 buffer. Conjugated bile acids (**Figure S6**) were solubilized in DMSO to a final concentration of 275 mM. A bile acid master mix was prepared by mixing all bile acids to a final concentration of 10 mM for each bile acid in Tris-HCl, pH 4.6 buffer. BSH concentrations were determined using a Bradford Assay and standardized to 1.03 μM +/- 0.27 μM. A 50 μL reaction was prepared with 1 mM of each amine, 1 mM of each bile acid, and approximately 0.4 μM of each BSH. Final concentrations were achieved with Tris-HCl, pH 4.6 buffer. The final pH was measured to be between 5-6 with pH paper. Reactions were incubated at 37 °C for 48 h and then quenched with methanol (1:3 reaction:methanol). Quenched reactions were centrifuged at 500 x *g* for 30 mins, and the supernatant was transferred to a flat bottom 96-well plate. Reaction products were stored at –80 °C until analysis was performed.

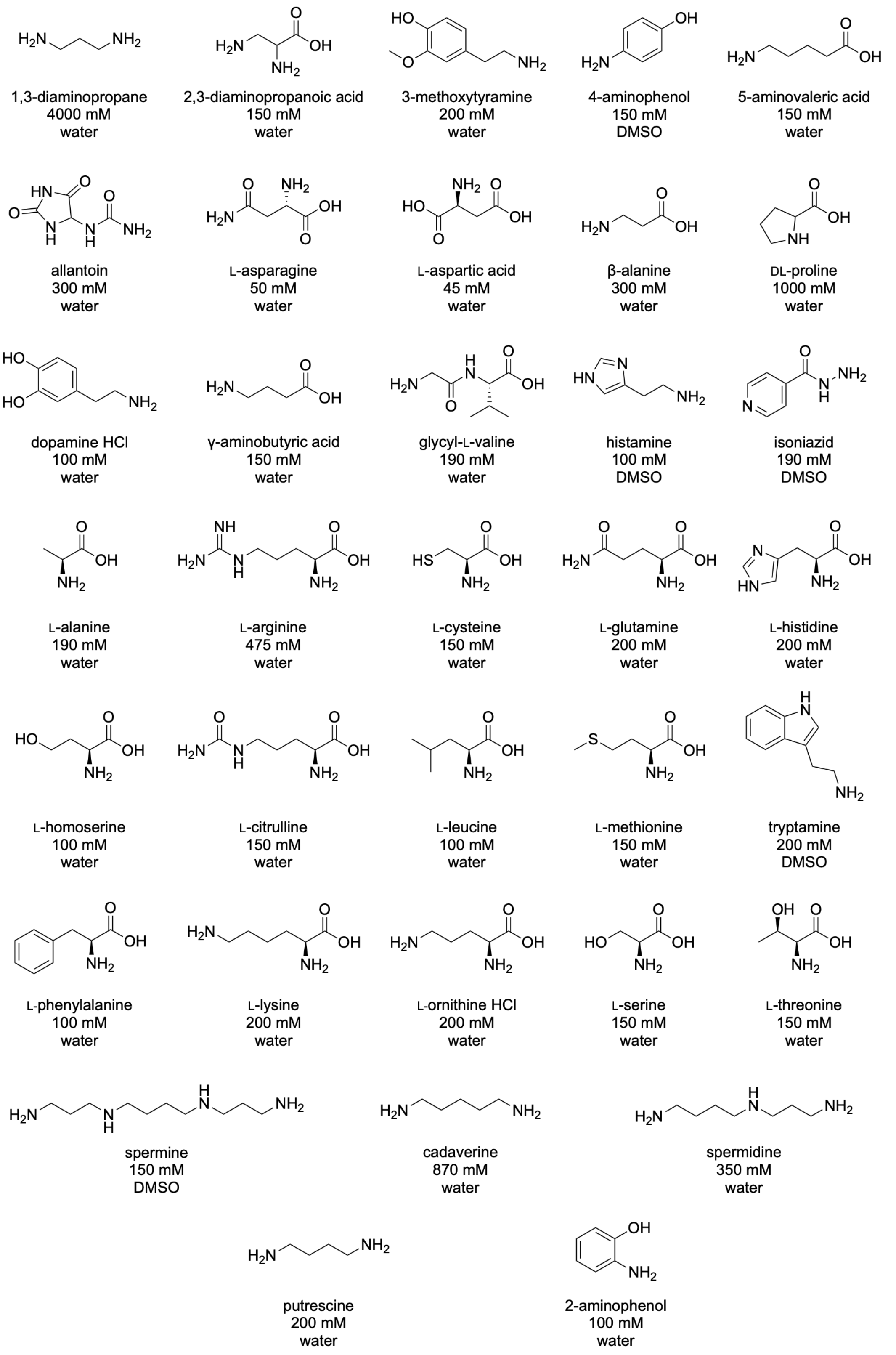

**Supplementary Figure S5. Amine substrates and preparation conditions.** All amines structures are shown with their name, preparation concentration (due to variable solubilities), and preparation solvent below.

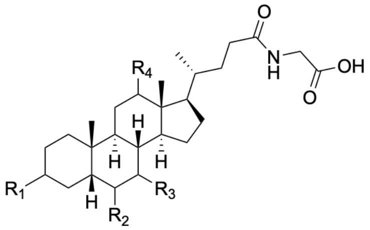

**Glycocholic acid substrates**

| | |
|---|---|
| **1** | $R_1 = R_3 = R_4$: β-OH; $R_2$: H |
| **2** | $R_1 = R_4$: β-OH; $R_2 = R_3$: H |
| **3** | $R_1 = R_3$: β-OH; $R_2 = R_4$: H |
| **4** | $R_1$: β-OH; $R_3$: α-OH; $R_2 = R_4$: H |
| **5** | $R_1 = R_2$: β-OH; $R_3 = R_4$: H |
| **6** | $R_1$: β-OH; $R_2 = R_3 = R_4$: H |
| **7** | $R_1$: α-OH; $R_2$: H; $R_3$: β-OH; $R_4$: =O |
| **8** | $R_1$: α-OH; $R_2$: H; $R_3$: α-OH; $R_4$: =O |
| **9** | $R_1$: α-OH; $R_2$: H; $R_3$: =O; $R_4$: β-OH |
| **10** | $R_1$: =O; $R_2$: H; $R_3$: α-OH; $R_4$: β-OH |

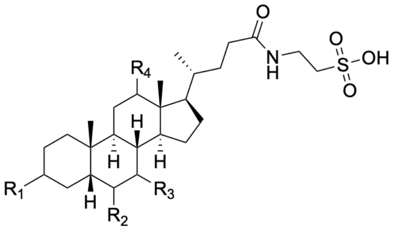

**Taurocholic acid substrates**

| | |
|---|---|
| **11** | $R_1 = R_3 = R_4$: β-OH; $R_2$: H |
| **12** | $R_1 = R_4$: β-OH; $R_2 = R_3$: H |
| **13** | $R_1 = R_3$: β-OH; $R_2 = R_4$: H |
| **14** | $R_1$: β-OH; $R_3$: α-OH; $R_2 = R_4$: H |
| **15** | $R_1 = R_2$: β-OH; $R_3 = R_4$: H |
| **16** | $R_1$: β-OH; $R_2 = R_3 = R_4$: H |
| **17** | $R_1$: α-OH; $R_2$: H; $R_3$: β-OH; $R_4$: =O |
| **18** | $R_1$: α-OH; $R_2$: H; $R_3$: α-OH; $R_4$: =O |
| **19** | $R_1 = R_3 = R_4$ : =O; $R_2$: H |

**Key**

| | |
|---|---|
| **1** | Glycocholic acid |
| **2** | Glycodeoxycholic acid |
| **3** | Glycochenodexoycholic acid |
| **4** | Glycoursodeoxycholic acid |
| **5** | Glycohyocholic acid |
| **6** | Glycolithocholic acid |
| **7** | 3α12k-glycocholic acid |
| **8** | 3α7α12k-glycocholic acid |
| **9** | 3α7k-glycocholic acid |
| **10** | 3k7α-glycocholic acid |
| **11** | Taurocholic acid |
| **12** | Taurodeoxycholic acid |
| **13** | Taurochenodeoxycholic acid |
| **14** | Tauroursodeoxycholic acid |
| **15** | Taurohyodeoxycholic acid |
| **16** | Taruolithocholic acid |
| **17** | 3α12k-taurocholic acid |
| **18** | 3α7α12k-taurocholic acid |
| **19** | 3α7α12k-taurocholic acid |

**Supplementary Figure S6. Conjugated bile acid substrates used in multiplexed assay.** Conjugated bile acids are grouped based on glycerol and taurine conjugation. Each number shows corresponding substitutions and a key provides the name for each compound. Each compound was prepared with DMSO as solvent to a concentration of 275 mM.

### IV. Untargeted LC-MS/MS Analysis

**LC-MS/MS data acquisition.** LC-MS/MS data were acquired on a Orbitrap Exploris 240 coupled to a Vanquish UHPLC system (Thermo Fisher Scientific). Chromatographic separation was performed on a Kinetex Polar C18 column (100 x 2.1 mm, 2.6 µm particle size, 100 Å pore size; Phenomenex), maintained at 40 °C. Mobile phases A and B consisted of $H_2O$ and acetonitrile, respectively, each containing 0.1% formic acid. The injection volume was 5 µL, and the flow rate was 500 µL per min. The LC gradient was programmed as follows: 5% B from 0 to 0.5 min; linear increases to 25% B at 1.0 min, 40% B at 7.5 min and 99% B at 8.5 min; and a hold at 99% B until 10.0 min. The column was returned to 5% B at 10.1 min and held until 11.1 min, followed by an additional wash at 99% B from 11.2 to 12.0 min. The mobile phase was returned to 5% B at 12.1 min and held until 13.0 min. Mass spectra were acquired using electrospray ionization in positive (ESI+) mode with spray voltage of 3.5 kV. Sheath, auxiliary, sweep gas settings were 50, 10, and 1 arbitrary units, respectively. The ion transfer tube and vaporizer temperatures were 325 °C and 350 °C, respectively. Full scan MS1 spectra were acquired over *m/z* 150-1500 at a resolution of 60,000 at *m/z* 200 with a RF lens of 70%, and automatic gain control (AGC) target set to standard with a maximum injection time of 100 ms. The MS/MS acquisition method was generated using the AcquireX Deep Scan workflow with pooled quality control (poolQC) samples to increase coverage. The resulting method was applied to all samples using top 10 data-dependent acquisition. The poolQC was injected every 10 samples. The precursor intensity threshold was 5e3, and the isolation window was 1 *m/z*. Precursors were fragmented using stepped collision energies of 25, 40, and 60 V. MS/MS spectra were acquired at a resolution 22,500 at *m/z* 200, with the first mass set to *m/z* 50, the AGC set to standard and the maximum injection time set to Auto. Spectra were recorded in profile mode with one microscan per spectrum. Precursors selected twice within 3 s were dynamically excluded for 5 s using a mass tolerance of ± 3 ppm, with isotope exclusion enabled.

**LC-MS/MS data processing.** The raw MS/MS spectra were converted to mzML files using MSconvert (ProteoWizard)[66] and deposited in GNPS/MassIVE with accession ID: MSV000097665. Feature extraction was performed using MZmine 4.2.0 with the following parameters: for mass detection, the factor of lowest signal was 5 for MS1 and 2.5 for MS2. For chromatogram building, the mass tolerance was set as 0.002 m/z or 10 ppm, the minimum consecutive scans as 4, and the minimum height as 5E4. Local minimum search for chromatographic deconvolution was performed with a minimum search range of 0.05 min, minimum ratio of peak top to edge of 1.8, and maximum peak duration of 1.2 min. The peaks were de-isotoped within 5 ppm *m/z* and 0.05 minutes retention time tolerances, aligned with 0.0015 *m/z* or 5 ppm mass tolerance and 0.2 minutes retention time tolerance, then gap-filled with 5 ppm *m/z* tolerance and 0.2 minutes retention time tolerance. Ion forms were assigned using the metaCorrelate module with a retention time tolerance of 0.02 min, minimum feature height of 3E4 and intensity threshold for correlation at 1E4. Feature height correlation was selected using Pearson similarity measure with a minimum correlation of 70 % across a minimum of 3 samples. The feature list was then exported as a feature quantification table (.csv) and an MGF spectra file. Metabolite annotations were performed with the default GNPS Spectral Libraries and the GNPS-BILE-ACID-MODIFICATIONS, GNPS-MASSQL-BILE-ACID-ISOMER and the NEW_MULTIPLEXED_FILTERED_LIB libraries using the Feature-Based Molecular Networking (FBMN) workflow in GNPS2. The spectra were filtered by removing MS/MS fragments within ±17 Da of the precursor *m/z* and to only keep the six most intense fragments in a ±50 Da window. The spectra were searched against the GNPS2 libraries with precursor and fragment ion mass tolerances of 0.02 Da and 0.05 Da, respectively, with a cosine score threshold of 0.6 and a minimum of 5 matched peaks. FBMN GNPS2 job link: https://gnps2.org/status?task=6af4031f410d4284b964b0d3faf5155b

**Selection of candidate bile acid amidates (BBAs).** To identify candidate BBAs, we first subsetted MS/MS spectra that had a spectral library match to entries in either the GNPS-BILE-ACID-MODIFICATIONS or GNPS-MASSQL-BILE-ACID-ISOMER libraries, which are the two bile acid-specific libraries. However, these spectral matches can include false-positive annotations as well as multiply modified bile acids. Because the present study focused specifically on single modifications at the C24 carboxyl group of bile acids, such spectra were excluded from further analysis. Therefore, the resulting candidate MS/MS spectra were then subjected to a second-stage MassQL filtering workflow. Stage 2 MassQL queries were adapted from our previously described workflow[11] and required both bile acid diagnostic fragment ions and detection of the expected protonated amine/amino-acid fragment generated upon cleavage of the bile acid-amine conjugate. This additional criterion was used to enrich candidate bile amidates modified at the C24 carboxyl group. Spectra satisfying both the bile acid diagnostic-ion criteria and the expected amine-associated mass-difference criteria were retained as putative BBAs. Using this workflow, 58 of 527 candidate MS/MS spectra were retained for downstream annotation and analysis.

Stage 2 MassQL queries used are below:

*Monohydroxy: QUERY scaninfo(MS2DATA) WHERE*
*MS2PROD=341.28:TOLERANCEMZ=0.01:INTENSITYPERCENT=5 AND*
*MS2PROD=323.27:TOLERANCEMZ=0.01:INTENSITYPERCENT=5 AND*
*MS2PREC=X AND MS2PROD=X-358.287:TOLERANCEMZ=0.01:INTENSITYPERCENT=5*

*Dihydroxy: QUERY scaninfo(MS2DATA) WHERE*
*MS2PROD=339.27:TOLERANCEMZ=0.01:INTENSITYPERCENT=5 AND*
*MS2PROD=321.26:TOLERANCEMZ=0.01:INTENSITYPERCENT=5 AND*
*MS2PREC=X AND MS2PROD=X-374.282:TOLERANCEMZ=0.01:INTENSITYPERCENT=5*

*Trihydroxy: QUERY scaninfo(MS2DATA) WHERE*
*MS2PROD=337.25:TOLERANCEMZ=0.01:INTENSITYPERCENT=5 AND*
*MS2PROD=319.24:TOLERANCEMZ=0.01:INTENSITYPERCENT=5 AND*
*MS2PREC=X AND MS2PROD=X-390.277:TOLERANCEMZ=0.01:INTENSITYPERCENT=5*

*Tetrahydroxy: QUERY scaninfo(MS2DATA) WHERE*
*MS2PROD=335.24:TOLERANCEMZ=0.01:INTENSITYPERCENT=5 AND*
*MS2PROD=317.23:TOLERANCEMZ=0.01:INTENSITYPERCENT=5 AND*
*MS2PREC=X AND MS2PROD=X-406.272:TOLERANCEMZ=0.01:INTENSITYPERCENT=5*

*Pentahydroxy: QUERY scaninfo(MS2DATA) WHERE*
*MS2PROD=333.22:TOLERANCEMZ=0.01:INTENSITYPERCENT=5 AND*
*MS2PROD=315.21:TOLERANCEMZ=0.01:INTENSITYPERCENT=5 AND*
*MS2PREC=X AND MS2PROD=X-422.267:TOLERANCEMZ=0.01:INTENSITYPERCENT=5*

**Repository-scale spectral searching using fastMASST.** Batch searches of the BBA reference MS/MS spectra were performed against publicly available tandem mass spectrometry data in the GNPS2 repository using fastMASST[57]. For each query spectrum, precursor and fragment ion mass tolerances were set to 0.05 Da, the minimum cosine similarity threshold was set to 0.70, and at least three matching fragment ions were required for a spectral match. fastMASST searches were used to identify spectra in public datasets with fragmentation patterns similar to the enzymatically generated BBA reference spectra. Search results were returned as Universal Spectrum Identifiers (USIs)[67], enabling individual matched spectra to be traced to their originating datasets and samples. These USIs were subsequently mapped to harmonized sample metadata in panReDU for downstream analysis of disease associations, and life-stage associations.

**Mapping BBA spectral matches to public metabolomics metadata.** fastMASST searches returned USIs corresponding to MS/MS spectra in public metabolomics datasets that matched the query BBA spectra. To provide biological context for these matches, we mapped their distribution across body parts, disease states, and life stages using an R-based workflow adapted from previously described repository-scale metadata analysis methods[68]. For each matched USI, sample-level metadata were retrieved from panReDU[58], a harmonized metadata resource for public metabolomics datasets. The panReDU metadata repository was downloaded and queried to identify the experimental condition, sample type, disease annotation, life-stage annotation, and corresponding sample identifiers associated with each USI. To account for differences in the number of publicly available samples across phenotypic categories, prevalence was normalized in two steps. First, for each BBA and each phenotype category, the number of matched files was divided by the total number of files in panReDU annotated with that phenotype. For example, the number of files containing a match to a given BBA in inflammatory bowel disease (IBD) samples was divided by the total number of IBD-associated files represented in panReDU. This yielded a phenotype-specific prevalence estimate for each BBA. Second, to compare the relative distribution of each BBA across the phenotypes in which it was detected, the phenotype-specific prevalence values were normalized to sum to 100% for each BBA. Thus, the resulting values represent the relative strength of association of each BBA with the disease states or life stages in which it was observed, after accounting for differences in dataset representation. These normalized values were visualized as heatmaps. Accompanying bar plots show the total number of metadata-associated files in which each BBA was detected.

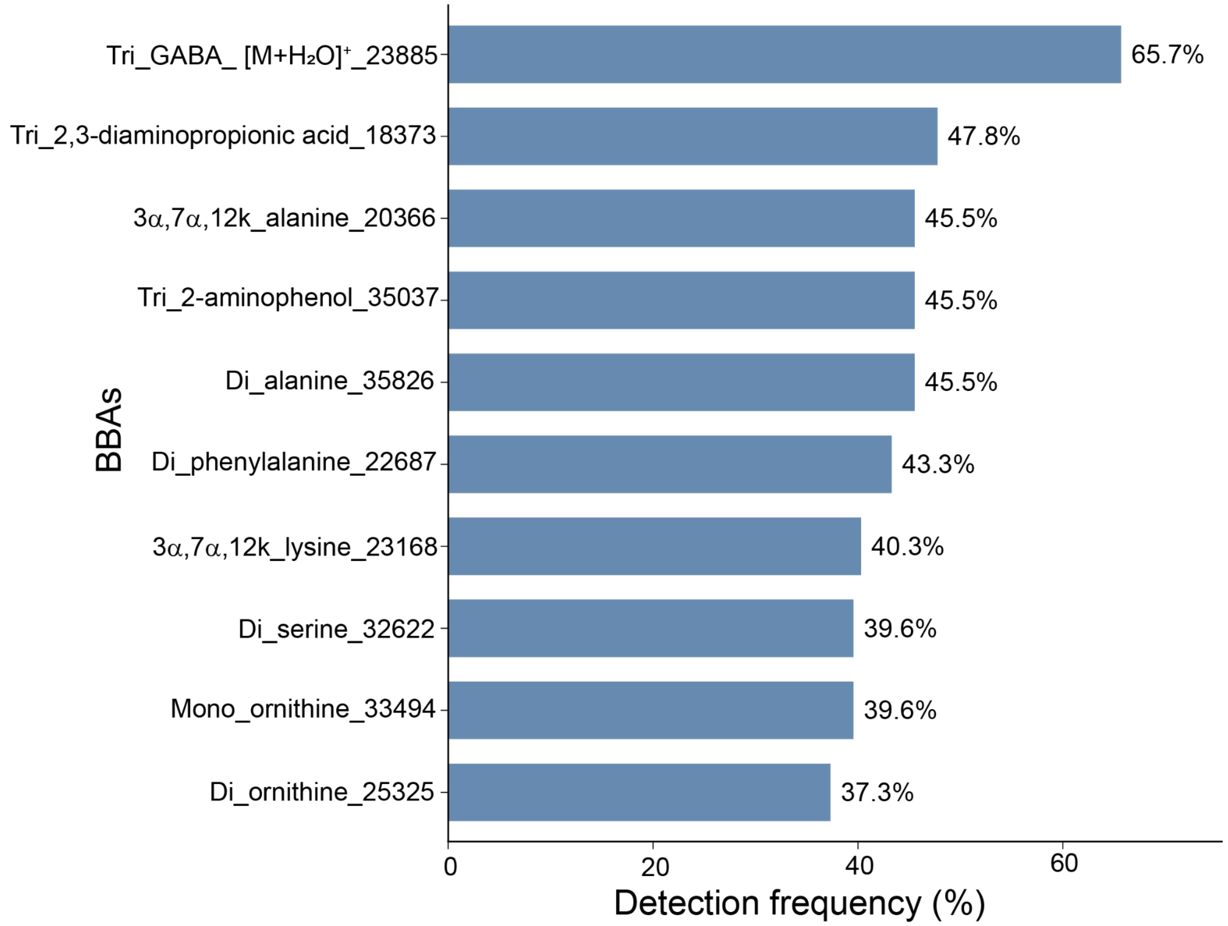

**Supplementary Figure S7. Frequency of the most commonly detected BBAs across the enzyme screen.** Bar plot showing the top 10 BBAAs ranked by detection frequency. Detection frequency was calculated as the percentage of screened enzymes for which a given BBAA was detected in at least one replicate. Percentages are indicated at the end of each bar.

### V. Proteomics

**Samples denaturation, reduction, and alkylation.** Recombinant proteins were mixed with 250 µL of Lysis buffer composed of 6M urea [Fisher Chemicals, CAT# U15-500], 7% SDS [Fisher BioReagents, CAT# BP166-500], 50 mM triethylammonium bicarbonate (TEAB) [Sigma-Aldrich, CAT# T7408-500ML], pH 7.7; supplemented with Thermo Scientific™ HALT™ Protease and Phosphatase Inhibitor Cocktail, EDTA-free (100X) [Fisher Scientific, CAT# PI78445] at a final concentration 1X. Samples were incubated for 20 minutes in a MixMate® Mixer [Eppendorf, CAT# 5353] at 1000 rpm, room temperature, and sonicated for 5 minutes in a Branson 2800 Ultrasonic Bath [Branson Ultrasonics, CAT# CPX952219R]. Next, 250 µL of each sample was transferred into a 2 mL deep well plate [Eppendorf, CAT# 30504305], and proteins were chemically reduced by the addition of 2.5 µL of 0.5 M dithiothreitol (DTT) [Invitrogen, CAT# 15508-013] and a 30-minute incubation at 47℃. Samples were cooled on ice for 5 minutes, and proteins were alkylated by the addition of 7.5 µL of 0.5 M iodoacetamide [Sigma-Aldrich, CAT# I1149] and a 45-minute incubation at room temperature in the dark. The alkylation reaction was stopped by adding 2.5 µL of 0.5 M DTT and incubating for 15 minutes at room temperature.

**Protein digestion.** Samples were acidified by the addition of 27 µL of 12% phosphoric acid [Sigma-Aldrich, CAT# 49685-500ML]), mixed with 1.5 mL of Binding buffer composed of 90% methanol [Fisher Chemicals, CAT# A456-4], 50 mM TEAB, pH 7.7; and loaded, ~350 µL at a time, onto 96-well S-Trap™ plate [ProtiFi, CAT# C02–96well-10], placed on a 1.1 mL deep well plate [Axygene, CAT# P-DW-11-C], by 1–2-minute centrifugation at 1500×*g*, room temperature. Wells were washed 5 times with 200 µL of Binding buffer by 1-minute centrifugation at 1500×*g*, room temperature. The S-trap plate was next centrifuged for 5 minutes at 1500×*g*, room temperature, to remove residual methanol, and transferred onto new 1.1 mL deep well plates. Trypsin digestion mix, composed of 5 µL (2.5 µg) of Sequencing Grade Modified Trypsin [Promega, CAT# V5113] and 120 µL of 50 mM TEAB, was loaded into each well, the plate was briefly centrifugated (~4 seconds at <1,500 x *g*, room temperature), the resulting flow-through was reapplied onto the wells, and the plate was incubated overnight at 37 ˚C. Peptides were eluted from the wells with 125 µL of 50 mM TEAB (1 minute centrifugation at 2000 x *g*, room temperature), 125 µL of 5% formic acid [Fisher Chemicals, CAT# A117-50] (1-minute centrifugation at 2,000 x *g*, room temperature), and 125 µL of 50% acetonitrile [Fisher Chemicals, CAT# A955-500] (3 minute centrifugation at 2000 x *g*, room temperature). S-Trap plate was discarded, 1.1 mL deep well plates containing eluted peptides were covered with a silicone sealing mat [Axygen, CAT# AM-2ML-RD], and samples were transferred to –80℃. Frozen peptides were dried using a Savant SPD111V SpeedVac Concentrator [Thermo Scientific, CAT# SPD111V-115] in line with Savant RVT5105 Refrigerated Vapor Trap [Thermo Scientific, CAT# RVT5105-115].

**Peptide desalting.** Dried peptides were suspended in 500 µL of 0.1% trifluoroacetic acid (TFA) [Thermo Fisher Scientific, CAT# 28901] and incubated for 20 minutes in a MixMate® Mixer [Eppendorf, CAT# 5353] at 1000 rpm, room temperature. Sep-Pak C18 96-well plate (100 mg sorbent) [Waters, CAT# WAT-186002321], placed on a 1.1 mL deep well plate, was washed with 1 mL of 100% acetonitrile and twice with 1 mL of 0.1% TFA by 1-minute centrifugation at 300 x *g*, room temperature. Samples were applied onto the wells and centrifuged for 1 minute at 300 x *g*, room temperature. Wells were washed five times with 1 mL of 0.1% TFA by 1-minute centrifugation at 300 x *g*, room temperature, and a Sep-Pak C18 96-well plate was transferred onto a new 1.1 mL deep well plate. Peptides were eluted with 750 µL of 40% acetonitrile, 0.5% acetic acid [Fisher Chemicals, CAT# A35-500] and 750 µL of 80% acetonitrile, 0.5% acetic acid by 1-minute centrifugation at 300 x *g*, room temperature.

Eluted peptides were transferred into a 2 mL deep well plate, covered with a silicone sealing mat [Axygen, CAT# AM-2ML-SQ], and transferred to –80℃. Frozen samples were dried using a Savant SPD111V SpeedVac Concentrator in line with a Savant RVT5105 Refrigerated Vapor Trap.

**Peptide quantification.** Dried peptides were suspended in 500 µL of 50% acetonitrile and incubated for 20 minutes in a MixMate® Mixer at 1000 rpm, room temperature. Pierce Quantitative Colorimetric Peptide Assay kit [Thermo Fisher Scientific, CAT# 23275] was used according to the manufacturer's recommendations to determine peptide concentration. Transferred 1 µg aliquots of each sample into 1.5 mL Protein LoBind Tubes, frozen at –80℃, and dried using Savant SPD111V SpeedVac Concentrator in line with Savant RVT5105 Refrigerated Vapor Trap.

**Sample reconstitution for mass spectrometry analysis.** Dried samples were suspended in 10 µL of 5% acetonitrile and 5% formic acid and mixed for 20 minutes in a Vortex-Genie 2 vortex [Scientific Industries, CAT# 00-SI-0236], sonicated for 5 minutes in a Branson 2800 Ultrasonic Bath, and centrifuged for 2 minutes at 16,000 x *g*, room temperature. Supernatants were transferred into SureSTART™ 0.2 mL Amber TPX vials with Glass Insert [Thermo Scientific, CAT# 6PK1655].

**Proteomics Mass Spectrometry Data Acquisition.** Proteomic spectral data were collected using a data-independent acquisition (DIA) approach with a Thermo Scientific Orbitrap Astral Mass Spectrometer. A 2.5 µL (250 ng) of each sample was separated by reverse-phase high pH liquid chromatography using Vanquish™ Neo UHPLC System [Thermo Fisher Scientific] and Aurora Elite™ XT 15×75 C18 UHPLC column [IonOpticks, CAT# AUR4-15075C18-XT]. Solvent A of the mobile phase consisted of 0.1% formic acid in water, while solvent B consisted of 0.1 % formic acid in 80 % acetonitrile. Samples were separated by a 25-minute method composed of a linear gradient of 1-4 % solvent B at minute 0-0.1 with a flow rate of 700 nL/min, a linear gradient of 4-12 % solvent B at minute 0.1-1.9 with a flow rate of 700 nL/min, a linear gradient of 12-22.5 % solvent B at minute 2-12 with a flow rate of 700 nL/min, a linear gradient of 22.5-40 % solvent B at minute 12-19.5 with a flow rate of 500 nL/min, a linear gradient of 40-99 % solvent B at minute 19.5-22 with a flow rate of 500 nL/min, and a isocratic flow of 99 % solvent B at minute 22-25 with a flow rate of 600 nL/min.

Data was acquired with the following settings: Experiment #1 [MS] Master Scan using Full scan at Orbitrap Resolution 240,000, scan range of 380-980 *m/z*, RF Lens 40 %, Custom AGC Target, Normalized AGC target 500%, Absolute AGC Value 5.000e6, Maximum Injection Time 3 ms, Microscans set to 1, Data type set to Profile, Positive polarity; Experiment #1 [DIA] Master Scan using DIA at Precursor Mass Range 380-980, DIA Window Type set to Auto, Isolation Window 2 *m/z*, Window overlap set to 0 *m/z*, Window Placement Optimization turned On, Number of Scan events set to 299, DIA Window Mode set to *m/z* Range, Normalized Collision Energy Type, HCD Collision Energy set to 25%, Astral detector type, scan range of 150-2000 m/z, RF Lens 40%, Custom AGC Target, Normalized AGC target 500%, Absolute AGC Value 5.000e4, Maximum Injection Time 3 ms, Microscans set to 1, Data type set to Centroid, Positive polarity, Loop Control set to Time with 0.6 sec.

**Proteomics Mass Spectrometry Data Search.** For the DIA analysis, an *in silico* predicted spectral library was generated from a UniProt reference proteome of *Escherichia coli* (strain B / BL21-DE3) (accession UP000001509) combined with sequences of 125 selected microbiome proteins (Supplementary table 1), using DIA-NN software (version 2.2 Academia) with the following settings: FASTA digest for library-free search/library generation; Deep learning-based spectra, RTs and IMs prediction; Protease = Trypsin/P; Missed cleavages = 1, maximum number of variable modifications = 1, and modifications = N-term M excision, C carbamidomethylation. Thermo Finnigan RAW spectra files with DIA data were imported into the DIA-NN software and searched against the generated spectral library using the following settings: Protease = Trypsin/P; Missed cleavages = 1; maximum number of variable modifications = 1; modifications = N-term M excision, C carbamidomethylation, Ox(M); use isotopologues; MBR; Heuristic Protein inference, No shared spectra; Protein interference = Genes; Neural Network classifier = Single-pass mode; Quantification strategy = Robust LC (high precision); Cross-run normalization = RT-dependent; Library generation = Smart profiling; Precursor FDR (%) = 1.0; Mass accuracy, MS1 accuracy, and Scan Window were used at the default (0) setting.

**Data and Code availability:** All proteomics data (.raw file and metadata) are publicly available online at MassIVE Repository (https://massive.ucsd.edu) under identifier: MSV000103172. Untargeted LC-MS/MS data is deposited to GNPS/MassIVE under the accession MSV000097665. Code for Figure 3 and Supplementary Figure S7 can be accessed here: https://github.com/mohantyipsita/2026_BSH_substrate_screening. Code for Figure 4 can be accessed here: https://github.com/ssp5682-create/BSH_study_code.
